# Deviation from baseline mutation burden provides powerful and robust rare-variants association test for complex diseases

**DOI:** 10.1101/2020.07.04.186619

**Authors:** Lin Jiang, Hui Jiang, Sheng Dai, Ying Chen, Youqiang Song, Clara Sze-Man Tang, Binbin Wang, Maria-Mercedes Garcia-Barcelo, Paul Tam, Stacey S. Cherny, Pak Chung Sham, Miaoxin Li

## Abstract

The identification of rare variants that contribute to complex diseases is challenging due to low statistical power. Here we propose a novel and powerful rare variants association test based on the deviation of the observed mutational burden in a genomic region from a baseline mutation burden predicted by weighted recursive truncated negative-binomial regression (RUNNER) on genomic features available from public data. Simulation studies show that RUNNER is substantially more powerful than state-of-the-art rare variant association methods (including SKAT, CMC and KBAC), while maintaining correct type 1 error rates under population stratification and in small samples. Applied to real data, RUNNER “rediscovered” known genes of Hirschsprung disease missed by current methods, and detected promising new candidate genes, including *NXPE4* for Hirschsprung disease and *CXCL16* for Alzheimer’s disease. The proposed approach provides a powerful and robust method to identify rare risk variants for complex diseases.

## Introduction

Rare genetic variants, typically defined as variants with frequency under 1% in the population, are recognized to play an important role in the development of complex genetic diseases^1-3^, which may explain a nontrivial fraction of “missing heritability”^4^ from genome-wide association studies. Recent advances in high throughput sequencing (HTS) technologies have made feasible genome-wide rare variants association studies on large samples feasible. For example, the UK10K project has uncovered highly penetrant rare variants for cardiometabolic traits 1% ^5^, and the Trans-Omics for Precision Medicine (TopMed) project will examine contributions of rare variants to heart, lung, blood, and sleep disorders^6^. Rare variants in the *TREM2* and *APP* genes have been associated with Alzheimer’s disease, and similar rare susceptibility variants have been reported for type 2 diabetes, myocardial infarction, osteoporosis, and other diseases^3^. Compared to common variants, rare variants are more likely to be recent and deleterious, and might more easily lead to the discovery of new biological mechanisms or drug targets, such as the lipoprotein pathway gene PCSK9 for low-density lipoprotein cholesterol levels^7^.

Despite these early promising findings, studies on rare susceptibility variants generally suffer from low statistical power^2^. As testing association for single rare variants have very low power, multiple-variants based association methods have been proposed^2^, which test for association between phenotype and multiple rare variants within a gene or genomic region. Multi-variant association is typically evaluated by comparing mutation burden between patients and controls, in terms of the mean ^8; 9^, and/or the variance ^10^. Despite the gain in power over single-variant tests, existing multi-variant association tests still require very large samples^11^ which are difficult and costly to obtain. To increase study efficiency, researchers have proposed the use publicly available population genomic data to boost the number of controls ^12; 13^. However, this may lead to the challenge of hidden population stratification resulting confounding and spurious associations, especially since it is more difficult to correct population stratification with rare variants than common variants ^14^.

Here, we propose a novel statistical method to detect rare susceptibility variants in genes or genomic regions. The proposed method used novel strategy for association testing, which involves modeling the baseline mutation burden across genes or regions conditional on various genomic features, and considering the deviation of the observed mutation burden against the predicted baseline. After characterizing the method’s type I error rate, power, and robustness to population stratification using simulated data, the method was applied it to real data sets to examine its power for detecting novel associations.

## Material and Methods

### <u>R</u>ecursive tr<u>u</u>ncated <u>ne</u>gative-binomial <u>r</u>egression (RUNNER)

Our proposed test is based on considering to observed rare mutation burden at a genomic region (typically a protein-coding gene) in cases, relative to its baseline expectation conditional on the characteristics of the genomic region (See workflow of the method in Figure 1d), estimated from a weighted recursive truncated negative-binomial regression (RUNNER). Truncated negative-binomial regression was chosen because of two features of the distribution of rare mutation counts in genes: inflated 0 mutation counts and over dispersion (Figure 1a and 1b). RUNNER was implemented in our high-throughput sequencing data analysis platform KGGSeq(V1.1+)^15; 16^ (http://grass.cgs.hku.hk/limx/kggseq/), fits a statistical model for the rare mutation counts of patients, and outputs standardized z-scores and p-values based on the excess of functionally-weighted rare mutations over their predicted baseline values.

**Figure 1:**
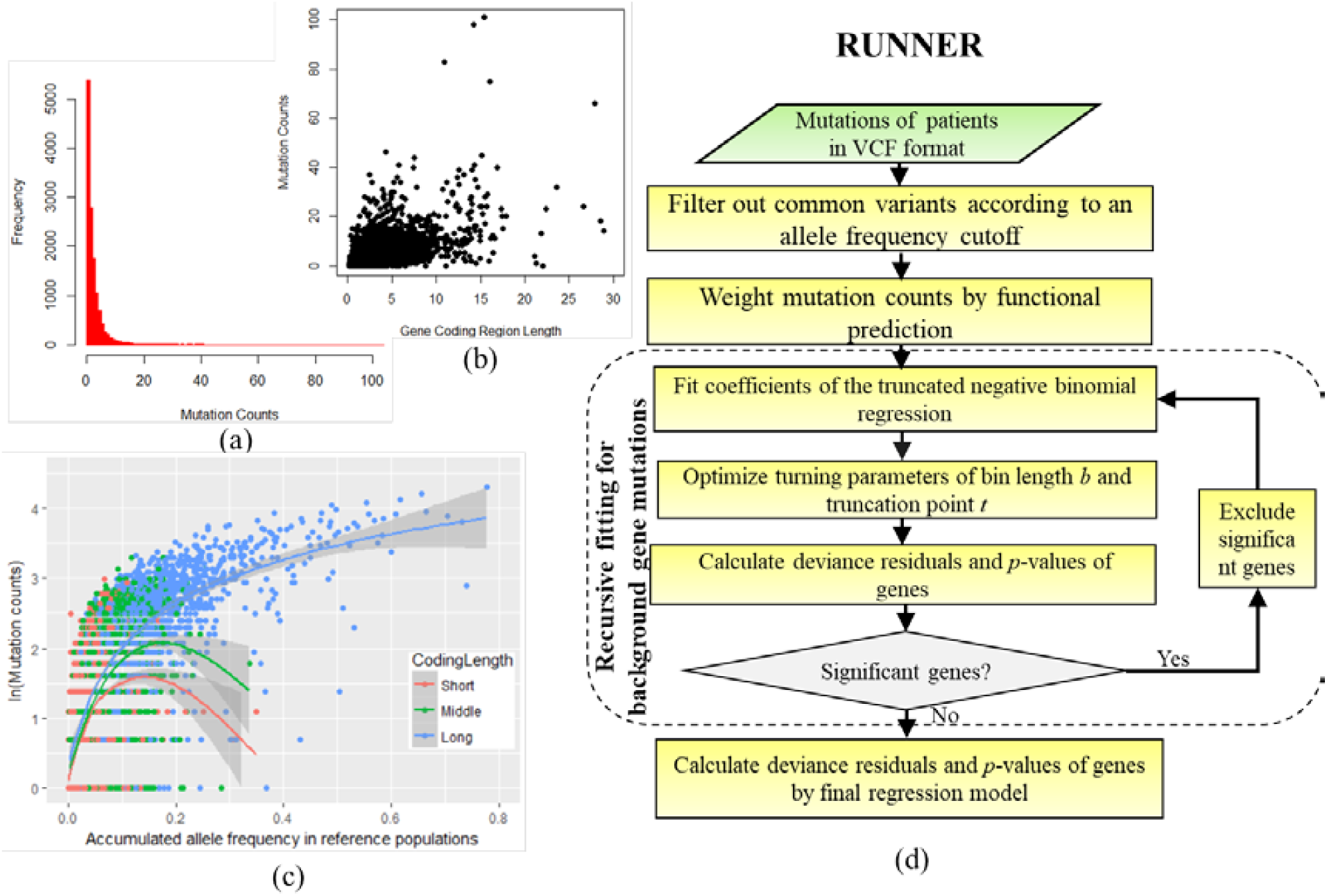
The distribution features of genes’ mutation counts and the proposed framework a): the histogram of rare mutation counts, b) the scatter plots of rare mutation counts of genes of different coding region length, c)the relationship between mutation counts and accumulated allele frequency in genes of short(<1.5kb), middle and long(>3kb) coding region length, d) The diagram of RUNNER procedure.

### The truncated negative-binomial regression model

With the assumption that a gene or region with excessive rare mutation burden in patients is more likely to confer disease risk, we extended the negative-binomial regression model to evaluate the burden for revealing potential susceptibility genes or loci of complex diseases. The burden is measured by excess of observed mutation counts relative to expected mutation counts. Let a gene or region *i* have *m*_*i*_ rare variants (including single nucleotide variants and InDels with allele frequency ≤ 1%) in sequenced patients of a disease after removing common variants defined in reference databases and removing variants which had allele frequency in patients of less than *d* (>1) folder in controls. Let a rare variant *j* of gene or region *i* has *n*_*i,j*_ mutant alleles among the patient samples (*n*_*i,j*_= 0,1,2, …). Assume the impact of mutant alleles can be further amplified by an integer weight *w*_*i,j*_, i.e., *c*_*i,j*_ = *n*_*i,j*_ * *w*_*i,j*_. The integer weight is converted from a bioinformatics prediction score (see details in the following section), *s*_*i,j*_ ∈ [0,1] by a ceiling function, 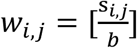, where *b* is a bin length parameter. According to our previous study on cancer somatic mutations ^17^, the total number of rare mutations in gene or region *i, y*_*i*_, can be assumed to approximately follow a negative binomial (NB) distribution:

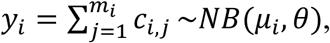

where *μ*_*i*_ is the expected number of mutations and *θ* is a dispersion parameter. The probability mass function (PMF) of NB distribution is 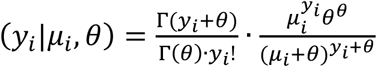, where Γ() is the gamma function and *y*_*i*_=0,1,2, ….

However, in a sample of typical size, many genes may have no rare mutations. That is, there would be an inflation of genes or regions with zero or low number of mutations(Figure 1a), which distorts the NB distribution. Therefore, we propose to model the mutation counts by a truncated negative binomial distribution. The PMF of the distribution with a truncated point parameter *t* is:

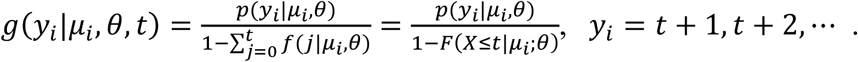

Meanwhile, the number of mutation counts of a gene is also affected by multiple factors. For instance, a long gene (say TTN, ∼305KB) tends to have more mutant alleles than a short gene (Say GJB1, ∼10KB). In the present study, we considered six predictive factors of the rare mutation counts at coding regions of genes,

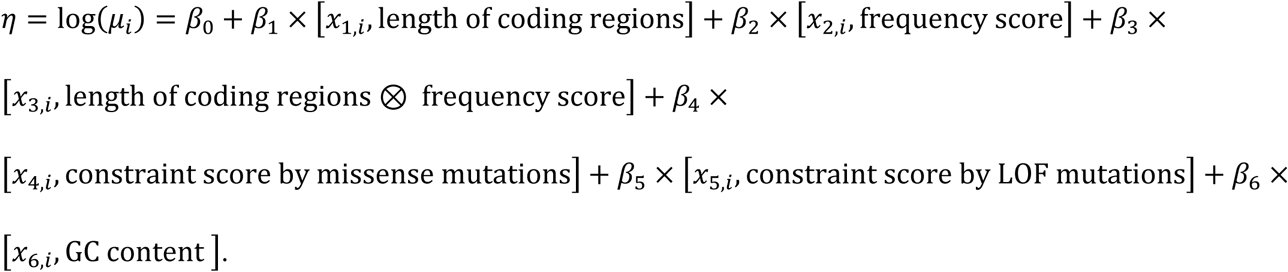

The frequency score, *x*_*2*_, is obtained from a reference database (e.g., 1000 Genomes Project) by matching allele-frequency and ancestry of the same region. The frequency score is produced by accumulating allelic frequencies of the same types of rare variants (see details below). The interaction factor, *x*_*3*_, is motived by the observation that genes of different coding region length may show different correlations between the mutation counts and allele frequencies (Figure 1c). The fourth and fifth predictors, constraint score which indicates a gene’s tolerance to missense and loss-of-function (LOF, e.g., protein-truncating) variants, are adopted from gnomAD ^18^. The OE_Miss and OE_LOF in the gene constrained file (https://gnomad.broadinstitute.org/downloads#gene-constraint) are used in the present study. The sixth predictor is the GC content of exon regions where the variants are located because mutation rate tends to be higher in GC-rich regions ^19^.

### Optimization of parameters

In the above model, the regression coefficients (*β*_0_, …, *β*_6_) and dispersion parameter (*θ*) are estimated by maximum likelihood approaches under the truncated negative binomial distribution. We derived the mathematical formulas of the first derivatives of the log-likelihood function, *l*(*y*_*i*_|*μ*_*i*_, *θ, t*) = ln[*g*(*y*_*i*_|*μ*_*i*_, *θ, t*)] = *l*_*i*_, for the maximum likelihood estimation (See details in the Supplementary Methods).

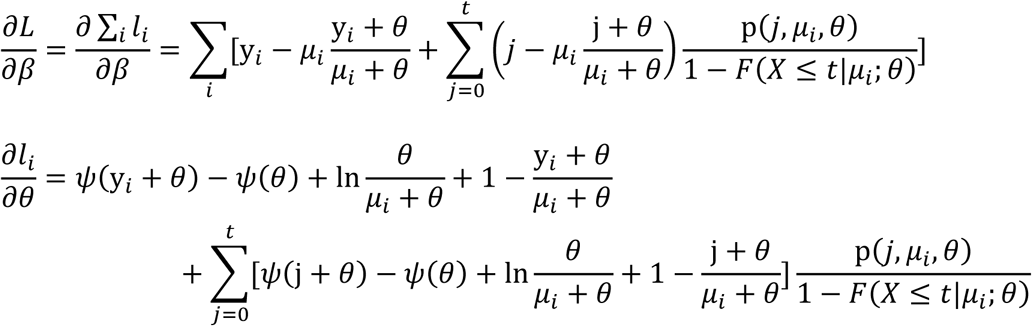

When fitting the six coefficients, the bin length *b* and truncation point *t* are set as constant. Given *b* and *t*, the fitted coefficients are used to estimate the expected mutation counts of gene *i*:

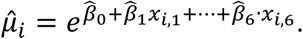

In the truncated negative binomial model, the deviance residual of the model at gene or region *i* is:

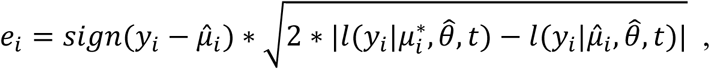

where *sign*(*x*) is the sign function of the raw residue, and the 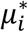 is the estimated mean given the observed count *y*_*i*_ and estimated 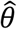 in a saturated model (See details in the Supplementary Methods).

The deviance residuals are further standardized by the estimated mean 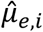 and standard deviation 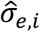 of the deviance residuals,

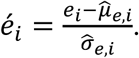

The standard normal distribution is used to approximate the corresponding p-value of *é*_*i*_:

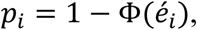

where *Φ*(*x*) is the cumulative distribution function of the standard normal distribution. Using simulated data, we showed that the truncation at negative binomial distribution led to approximately uniform distribution of the p-values (Figure S1). This is similar to the results in real data (Figure 2 and S2).

**Figure 2:**
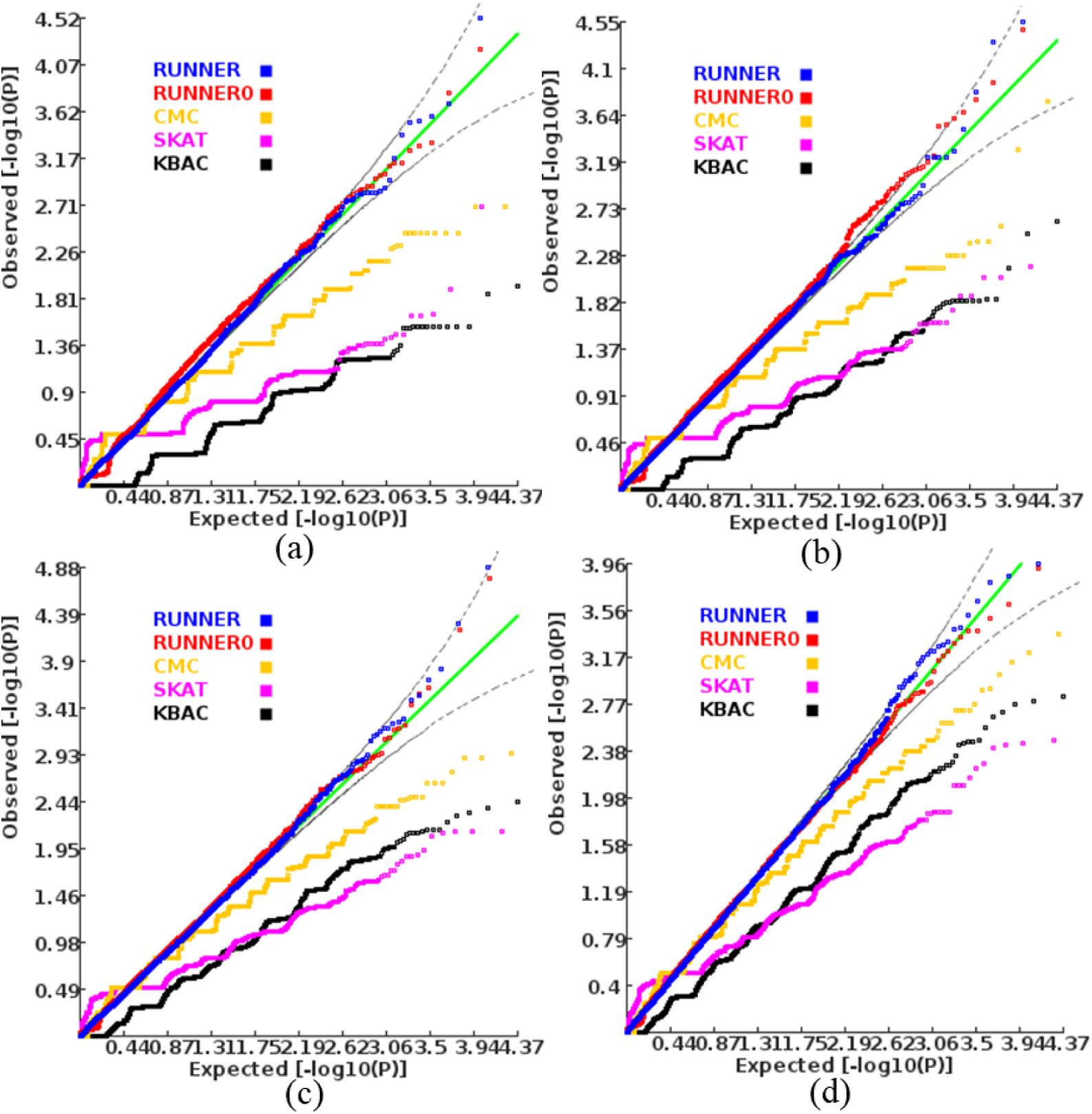
Quantile-quantile plot of p-values produced by different methods a) when sample size of cases is a)75; b) 100; c) 150; d) 200 and equal numbers of controls. RUNNER: the proposed approach; RUNNER: RUNNER0 using original mutation counts at variants. CMC, SKAT, and KBAC are three widely-used approaches for gene-based association tests with rare variants in case-control samples.

We built a grid-search procedure to explore the optimal values in their reasonable ranges. The bin length *b* is assumed to range from 0.05 to 1 and the truncation point *t* is assumed to range from 0 to 5. The search intervals of *b* and *t* are 0.05 and 1 respectively. Therefore, the grid-search has 120 (=20×6) *b*-*t* combinations in total. We adopted the mean log fold change (MLFC) ^20^ of p-values to quantify the departure from uniform distribution. The minimal MLFC is zero. A larger MLFC indicates a higher departure. The 120 combinations are ranked according to MLFC of potential background genes in ascending order and number of significant genes/regions (FDR *q*>0.05) in descending order respectively. The optimal *b* and *t* are defined as the one resulting in the minimal summation rank, which balances the uniform distribution departure of p-values and number of significant genes/regions.

### The recursive procedure

Note that the truncated negative binomial regression is proposed to model distribution of rare mutations at background genes or regions, i.e., the null hypotheses. The excessive mutations in true susceptibility genes or regions may harm the model fitting. Therefore, we further propose a recursive procedure to approximate the null hypothesis of gene mutations by removing significant and suggestively significant genes or regions. The following are the main steps of the recursive regression procedure (Also see the entire diagram in Figure 1d):

1. Perform the truncated negative binomial regression on all genes in a genome and calculate p-value of mutation burden for each gene by RUNNER.
2. Exclude significant genes according to a p-value cutoff, say, FDR of 0.2.
3. Re-perform the truncated negative binomial regression on the remaining genes.
4. Repeat Step 2 and 3 until no genes are excluded to obtain a converged model.
5. Calculate the deviance residues and p-values of all genes by the converged model.

#### Preparation of functional scores at mutations

As mutations are not equally important in terms of their impact on gene function, it is reasonable to use prior weights to amplify mutant alleles with higher functional impact. We adopted our previous ensemble score^15; 16^ from 19 prediction scores by a logistic regression model (See the names and descriptions of the tools in Table S6). The basic rationale of these predictions is that variants which are evolutionarily conserved and have large effects on physical-chemical properties and protein structure may have higher damaging effect on a protein of a gene^21^. A posterior probability of disease-causal potential is produced by combing 19 existing tools under a logistic regression framework. The posterior probability is further standardized into a range of (0, 1) according to their posterior probability. A variant with the largest posterior probability in the genome has the standardized ranking score of 1. A larger ranking score indicates a higher functional or pathogenic impact of the mutant allele on the gene’s function. For variants with missing scores, the average prediction score in the gene is used. The procedure of generating ranking scores has been implemented into KGGSeq for the truncated negative binomial regression.

#### Preparation of frequency scores

The above frequency score is created by accumulating allelic frequencies at variants after matching allele frequency, variant type and population ancestry. Define a gene or a region *i* which has *k* rare variants in a reference sample. A variant *j* of the gene or region *i* has allele frequency *f*_*i,j*_ and the above integer functional weight *w*_*i,j*_. The frequency score is then equal to:

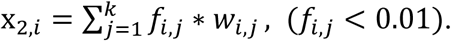

When there is more than one reference sample, the averaged allele frequencies of multiple samples is used for the frequency score. In the present paper, we adopted the allele frequencies of sequence variants from the 1000 Genomes Project and the exomes in Genome Aggregation Database (gnomAD, V2.1) as reference samples. Alternatively, one can also use the frequency scores calculated in other reference samples and even in local control samples. For the 1000 Genomes Project, the variant sites and allelic counts (Phase3 V6) of five different ancestry panels (African, Mixed American, East Asian, European, and South Asian) were downloaded from ftp://ftp.1000genomes.ebi.ac.uk/vol1/ftp/release/20130502/. For gnomAD, the variant sites and allelic frequencies of 7 ancestrally different panels (East Asian, South Asian, African/African American, Latino, Finnish, Non-Finnish European, and Ashkenazi Jewish) were downloaded from http://gnomad.broadinstitute.org/downloads. In gnomAD, we only included the variants and allelic frequencies from subjects labelled as controls. The variants were mapped onto genes according to physical positions in two gene models (RefGene and GENCODE) and annotated with gene features by KGGSeq^22^. The integer functional weight of these variants was derived from the logistic regression models ^15; 16^ as described above.

#### Alternative gene- or region-based association tests for rare mutations

We compared the performance of RUNNER with three widely-used association tests for rare mutations. The three methods used different model to evaluate the mutation burden between cases and controls.

- Combined Multivariate and Collapsing (CMC) method ^8^: The CMC method is a type of direct burden test for case-control samples. The variants were divided into multiple groups according to the allele frequencies. Within a group, mutations at multiple variants were collapsed. A multivariate test (Hotelling’s T2 test) was then employed to test whether the collapsed mutations of all groups were associated with case-control status.
- The sequence kernel association test (SKAT) ^10^: SKAT assumed the effects of rare variants on disease risk followed an arbitrary distribution with a mean of zero and an unknown variance. The variants without effect would have zero variance. A score-based variance-component test in a mixed model was then constructed to examine whether the variance was equal to zero.
- The kernel-based adaptive clustering model ^23^ (KBAC): KBAC considered genotypes of all rare variants in a gene or region as multi-site genotype vectors. The disease risk for a multi-site genotype was modeled using a mixture distribution. An adaptive weighting procedure was then used to produce weights of each genotype for their causality or susceptibility to a disease based on a known component called kernel.

The three methods were implemented into a user-friendly tool, RVTESTS^24^. Our high throughput sequencing data analysis platform, KGGSeq (http://grass.cgs.hku.hk/limx/kggseq/), has integrated RVTESTS as a third-party analysis component. On KGGSeq, we inputted the called sequence variants in VCF format and called RVTESTS for the gene-based association tests by the three approaches with default parameter settings in parallel.

#### Computer simulations investigate type I error and power

To keep the distribution of background rare mutations, we used a semi-simulation procedure to investigate the statistical type I error and power of RUNNER. In a database of 493 whole genome-sequenced subjects, which were used as healthy controls in a genetic study of Hirschsprung’s disease ^25^, we randomly drew *m* (=75, 100, 150 and 200) subjects to form a new sample. For the type I error, the RUNNER was used to calculate *p*-values of all genome-wide genes in the new samples. The quantile-quantile (QQ) plot of -log10 (p-value) under uniform distribution was used to check distribution of p-values, particularly at the tail of small p-values. In addition, the MLFC was also adopted to quantify the deviation of *p*-values from uniform distribution in general ^20^. A MLFC of > 0.3 indicated serious deviation from the null distribution ^20^. To investigate the statistical power, we made an artificial patient sample by randomly inserting several causal mutations (simulated according to a pre-set frequency) into genomes of some subjects of a drawn sample. Two genes (TCF4, and TIE1) were set as the targeted susceptibility genes in the simulations. Either gene was assumed to have 4-5 non-synonymous susceptibility variants (See the variants in Table S1). Each variant had an expected frequency of 1-1.5% at risk alleles in the artificial patient sample. The patient sample was analyzed by RUNNER to detect the genes with the rare risk mutations. In addition, we also randomly draw another batch of *m* subjects from the remaining whole-genome sequencing subjects and treated them as a matched control sample. The mutation rate of the variants in the assumed susceptibility variants was set as 10^−8^ in a control sample. The case-control samples were analyzed for power comparison. We produced *t* case-control samples for the association tests. Assume a test produced genome-wide significant p-values in *k* samples at a tested gene. The power was estimated as k/t at this gene.

In addition, we also used subjects from 1000 Genomes (1KG) Project to check how the methods were sensitive to population stratification. Two types of samples were set. The first type of sample contained 250 subjects from the CEU panel as cases, and 125 subjects from the remaining CEU panel and 125 subjects from the EAS panel as controls. The second type of samples included 500 subjects from the CEU panel as cases and 500 subjects from the EAS panel. The samples were analyzed by RUNNER and three alternative methods for association at rare variants. No genes were assumed to be related to the case/control membership, which was a mimic of null hypothesis. A significant association was spurious association due to population stratification

#### Real high-throughput sequencing datasets

Two real high-throughput sequencing datasets were used to validate the performance of the proposed method. Table S2 summarizes the disease names and sample information. All the sequencing samples were obtained with Institutional Review Board approval in either Hong Kong or in the mainland China. The short reads produced by Illumina HiSeq 2000 were mapped onto the reference genome (HG19) by BWA v0.7.17^26^. The redundant reads were removed by Picard (https://broadinstitute.github.io/picard/). The sequence variants were called by GATK v3.8 ^27^. A stringent quality control (QC) procedure at the variants was performed on KGGSeq V1.1 (http://grass.cgs.hku.hk/limx/kggseq/). The variants or genotypes failing to pass the following QC criteria were excluded: Hardy-Weinberg test p value ≤0.001, genotyping quality Score (Phred Quality Score) <20, read depth per genotype <8, having 3 or more alleles, genotyping rate <90% in a sample. Supplementary Table S2 also lists the initial and retained sequence variants in each dataset.

#### Detection of susceptibility genes with rare variants by the proposed methods and three alternative methods

All analyses for detecting susceptibility genes with rare variants were carried out on our KGGSeq platform (http://grass.cgs.hku.hk/limx/kggseq/). After the QC, common variants were then filtered out according to their alternative allele frequency (allele frequency > 0.01) in two databases, the 1000 Genomes Project and gnomAD exomes from East Asian panels. The variants were mapped onto genes according to two gene models (RefGene and GENCODE) and annotated with gene features (missense, synonymous, splicing, etc.). In the uncommon scenario that a variant belongs to multiple overlapped genes, the variants were assigned to each of the overlapped genes. Gene-based tests were then carried out to detect potential susceptibility genes of the diseases with the rare non-synonymous variants and splicing variants. For the gene-based association test by the three alternative methods [CMC ^8^, KBAC ^23^, and SKAT ^10^], KGGSeq produced all input of these tools and automatically launched RVTESTS for the analysis in parallel. For the baseline mutation burden test by RUNNER, KGGSeq directly ran its own codes to perform the analysis. KGGSeq predicted functional or pathogenic potential at a non-synonymous variant with a score (ranging from 0 to 1) and used the score to weight the mutant allele counts for RUNNER.

## Results

### Type I errors of the proposed and alternative methods for genome-wide association screening

To check whether RUNNER is valid for statistical inference, we first investigated its genome-wide statistical type I error for detecting susceptibility genes with rare mutations by random sampling approach. Similar to the simulated negative binomial data (Figure S1), the distribution of *p*-values produced by RUNNER for mutation counts in real samples was also very close to uniform distribution (in Figure 2). This is also true even when the sample size was as small as 75. As expected, in the special case of no weights at variants, RUNNER also generated approximately uniformly-distributed *p*-values. In contrast, the three widely-used gene-based association methods (CMC^8^, SKAT^10^, and KBAC^23^) for case-control analysis showed deflated *p*-value distribution. The time-consuming permutation (*n*=10^6^) SKAT, and KBAC did not rescue the deflation in the small samples. This was consistent with the consensus that existing statistical association tests for rare variants had generally invalid statistical properties in small or even moderate samples^28^. Relatively speaking, the p-values produced by CMC deviated from uniform distribution less than that produced by SKAT and KBAC. The deflation of the three methods became smaller as sample sizes increased. The valid type I error of RUNNER guarantees valid interpretation of RUNNER’s p-values for statistical inference.

### Robustness of RUNNER to population structure

We also found RUNNER was much less sensitive to population stratification in terms of the type 1 error inflation. In practice, it is very difficult to correct population structure for rare variants in association analysis. Fortunately, RUNNER seemed immune to population structure regardless of the degree. In the first type of sample in which 50% controls were from stratified populations, RUNNER showed no trend of inflation for baseline mutation burden test (Figure 3(a)). The unweighted version of RUNNER also showed uniformly distributed p-values. In contrast, the 50% of population stratification in controls resulted in very severe inflation of statistical significance at the three alternative association tests. There were over 200 genes with p-values <2.5E-6 by CMC and SKAT. KBAC had less inflation than CMC and SKAT but the trend of inflation was clear as well (Figure 3(a)). In another worse scenario, when all control subjects (*n*=500) had different ancestry from the cases (EAS vs. EUR), the inflation of statistical significance became even severer (Figure 3(b)). Hundreds of genes had p-values <2.5E-6 by CMC and SKAT. However, RUNNER still showed no inflation of statistical significance, in which the p-values followed uniform distribution approximately (Figure 3(b)). This analysis suggested RUNNER could be recommended for samples with large population stratification or batch effects as well.

**Figure 3:**
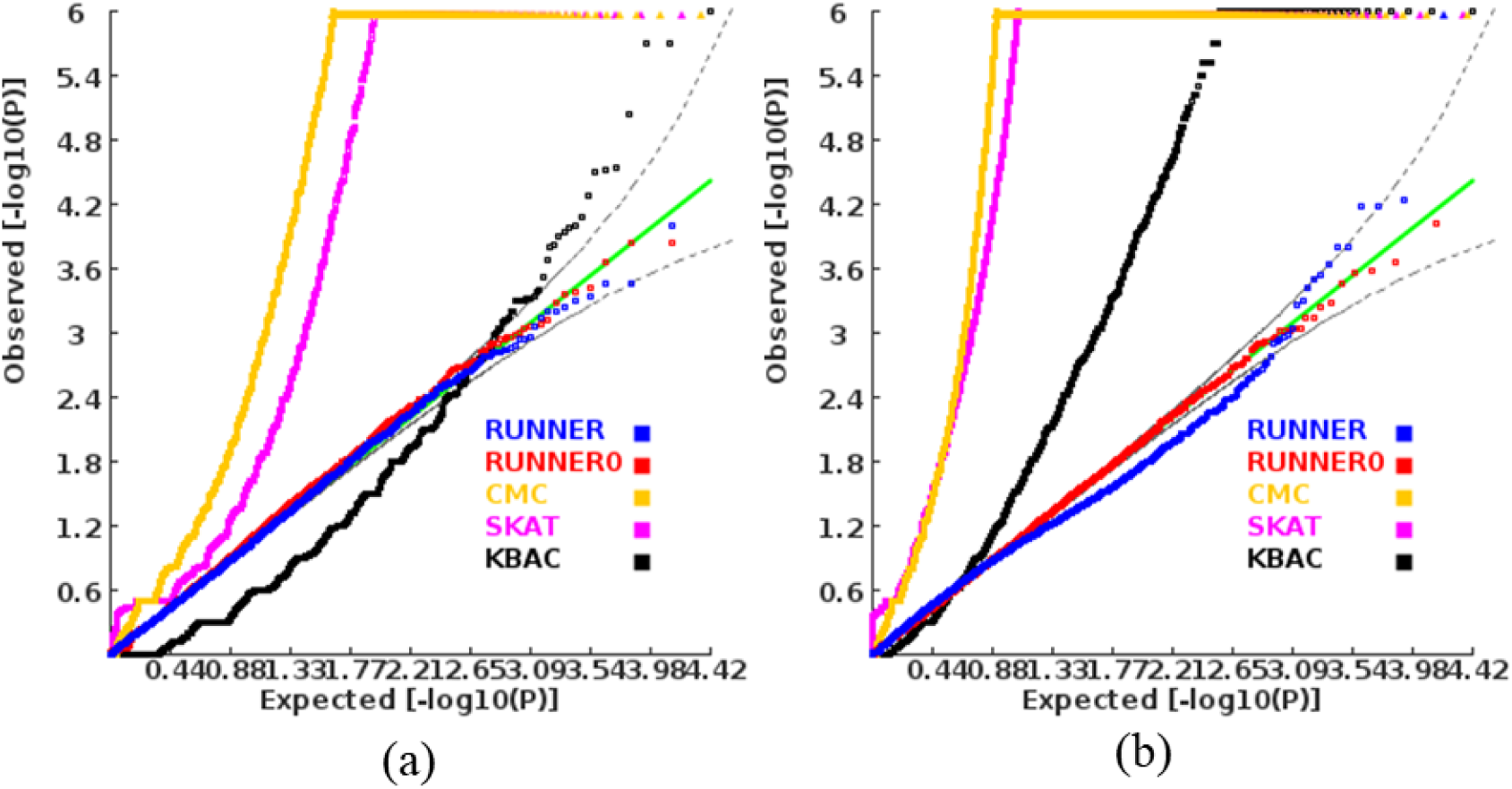
The p-value QQ plot of different tests in stratified population a) half of controls were from an ancestrally different population. b) all controls were from an ancestrally different population.

### The statistical power of the proposed and alternative methods for genome-wide association screening

We finally compared the statistical power of RUNNER with the three widely-used rare variant tests by semi-simulation analysis. The diseased samples were simulated by randomly inserting susceptibility alleles into real genomes of healthy subjects. Multiple rare missense mutations in TCF4 and TIE1 genes (See the mutation list in Table S1) were assumed in the simulation. TCF4 was repeatedly validated as a susceptibility gene for schizophrenia^29^ and TIE1 was suggested to play an important role in vascular development and pathogenesis of vascular diseases^30^. The RUNNER showed the highest power among all the compared tools (Figure 4). When the sample size was 100 patients and 100 controls, it achieved 58% and 68% power to detect TIE1 and TCF4, respectively, according to an exome-wide p-value cutoff 2.5E-6. The higher power at TCF4 than TIE1 was because TCF4 had more assumed causal mutations (5 vs. 4), shorter coding region (2.83 vs. 3.64), and lower accumulated allelic frequency in reference populations. When the sample size increased 50% (*n*=150), the power of RUNNER increased to 76% and 95% at the two genes, respectively. When no prior weights were imposed at the variants (the unweighted version of RUNNER), the power decreased. But it was still much more powerful than all the three alternative association tests for detecting both genes. When the sample size was 200 patients and 200 controls, the unweighted RUNNER achieved 87% power to detect TCF4. In contrast, the two gene-based association tests (SKAT and KBAC) had almost no power except that the CMC had 15% power to detect TIE1 in samples comprising 200 patients and 200 controls. This is consistent with previous studies that existing methods for rare variants had very low power in small or moderate samples^11^. Therefore, RUNNER achieved the best power to detect genes with rare susceptibility mutations based on its baseline mutation burden across genes.

**Figure 4:**
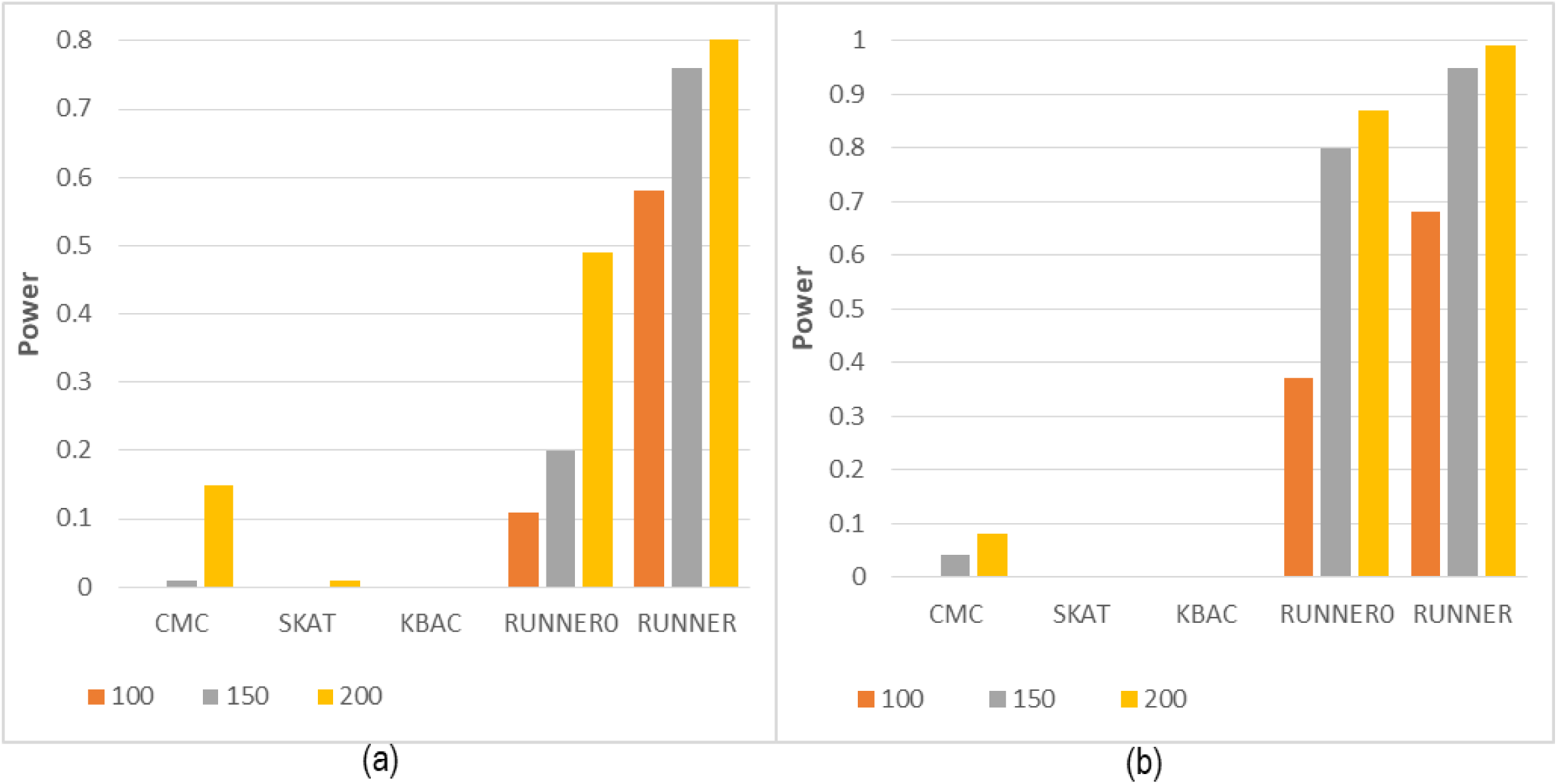
Performance comparison of different methods a) power of methods for detecting gene TCF4 b) power of methods for detecting TIE1 gene. For each scenario, 100 datasets were simulated. The assumed rare susceptibility variants can be seen in Table S1.

### Validation of RUNNER in real datasets of complex diseases

We further validated the performance of RUNNER at two different complex diseases (Hirschsprung disease and schizophrenia) with high throughput sequencing data (See summary description in Table S2). Genes were tested for rare mutation burden in patients by RUNNER. Besides, we also compared RUNNER’s results to three widely-used rare mutation association tests, CMC^8^, KBAC^23^, and SKAT^10^.

#### Hirschsprung disease

Hirschsprung disease (HSCR) is a highly heritable disorder with significant phenotypic heterogeneity and has a highest incidence rate among Asians (2.8/10,000 live births)^31^. Our dataset comprised 443 Asian HSCR patients and 493 Asian control individuals assayed by whole genome sequencing (The detailed sample information can be seen in the original paper^25^). The p-values of RUNNER, CMC and KBAC approximately followed the uniform distribution while SKAT showed deflation according to the Q-Q plots (Figure S2). Table 1 lists the top 10 genes detected by RUNNER. The well-known HSCR susceptibility gene, RET ^32; 33^, was obtained a significant p-value (*p*=3.03E-9) and prioritized as the top gene by RUNNER. In these cases, RET had 61 rare non-synonymous mutant alleles and its weighed allelic score was 122. The unweighted version of RUNNER also produced a highly significant p-value (*p* =2.48E-8) at RET and prioritized it as the top 1 gene. However, none of the alternative methods prioritized RET as the top 1 gene with the same set of rare variants. The association p-values by CMC, SKAT, and KBAC were 1.88E-05, 3.89E-04 and 7.0E-6 respectively, which did not survive the genome-wide significance level, 2.5E-6. The top 1 gene by CMC and KBAC was OGG1 (*p*=2.01E-6 and 1.0E-6) while SKAT ranked ZNF276 as the top 1 gene (*p*=2.97E-4). We failed to find literatures to support their contribution to the development of HSCR. A more remarkable instance can be seen in EDNRB, which is also a well-known HSCR susceptibility gene 33; 34. EDNRB was the 4^th^ most significant gene by RUNNER (*p*=4.67E-5). Unfortunately, CMC, SKAT and KBAC nearly neglected this gene, with a rank of 54, 199, and 58 respectively. The unweighted version of RUNNER only ranked it 39^th^ in all tested genes. This was because EDNRB only had 15 rare non-synonymous mutations. It was the weight that boosted its significance among all compared genes.

**Table 1:** Top 10 genes prioritized by RUNNER in Hirschsprung disease dataset

| Gene Symbol | Mutant alleles | Weighted alleles | Region Length | EAS | OE_LOF | OE_MIS | ExonGC | Residual<sub>0</sub> | P<sub>0</sub> | Residual | P |
| --- | --- | --- | --- | --- | --- | --- | --- | --- | --- | --- | --- |
| RET | 61 | 119 | 3.477 | 0.039 | 0.896 | 0.0391 | 0.585 | 5.452 | 2.48E-08 | 5.815 | 3.03E-09 |
| NXPE4 | 21 | 57 | 1.66 | 0.015 | 1.02 | 1.11 | 0.415 | 3.273 | 5.33E-04 | 4.913 | 4.49E-07 |
| CDCA4 | 19 | 45 | 0.731 | 0.021 | 0.934 | 0.452 | 0.531 | 3.419 | 3.14E-04 | 4.410 | 5.18E-06 |
| EDNRB | 15 | 36 | 1.797 | 0.011 | 0.806 | 0.329 | 0.410 | 2.680 | 3.68E-03 | 3.907 | 4.67E-05 |
| GAB4 | 33 | 50 | 2.105 | 0.015 | 1.15 | 0.815 | 0.578 | 3.645 | 1.34E-04 | 3.862 | 5.62E-05 |
| OGG1 | 26 | 48 | 1.732 | 0.018 | 1.1 | 0.885 | 0.576 | 3.342 | 4.16E-04 | 3.853 | 5.84E-05 |
| PHYKPL | 22 | 44 | 1.452 | 0.022 | 1.08 | 0.961 | 0.562 | 3.029 | 1.23E-03 | 3.612 | 1.52E-04 |
| PLOD2 | 20 | 41 | 2.484 | 0.022 | 0.921 | 0.439 | 0.366 | 3.096 | 9.82E-04 | 3.577 | 1.74E-04 |
| BECN1 | 15 | 29 | 1.408 | 0.009 | 0.666 | 0.151 | 0.481 | 3.041 | 1.18E-03 | 3.562 | 1.84E-04 |
| SEC24D | 20 | 50 | 3.235 | 0.032 | 0.83 | 0.405 | 0.436 | 2.513 | 5.99E-03 | 3.513 | 2.21E-04 |

Notes: Mutant alleles refer to rare non-synonymous mutant alleles. Region Length and ExonGC are the length and CG content in exon regions. Residual means deviance residue. Residual<sub>0</sub> and P<sub>0</sub> denote the residual and p-values of unweighted version of RUNNER.

Besides the two known susceptibility genes, RUNNER also detected another gene with significant rare mutation burden in the HSCR patients, NXPE4. The NXPE4 (*p*=4.49E-7), is a promising candidate gene. NXPE4 encodes neurexophilin and PC-esterase domain family member 4. Although it needs further investigation to reveal whether NXPE4’s dysfunction is related to congenital defect of the enteric nervous system in HSCR patients keeps unknown ^35^, there have been researches showing NXPE4 linked to several diseases originated in colons. NXPE4 is a prognostic biomarker for colorectal cancer^36^, and also associated with ulcerative colitis^37^. The high expression of NXPE4 had a significantly improved prognosis of colorectal cancer, regarding the overall survival of patients^38^. In addition, it has specifically high expression in colons (See details in database REZ, http://grass.cgs.hku.hk/limx/rez/).

In the fitted RUNNER model, all the six predictors significantly contributed to the prediction of mutation count at background genes. As expected, the region length and gene frequency scores were positively related to the rare mutation counts of genes (*p*<1.47E-164), which means longer coding regions and higher mutation frequency genes tend to have more rare non-synonymous mutations. Consistent with previous studies ^19^, genes with higher CG content also tended to have more rare non-synonymous mutations (*p*<1.33E-30). The gene’s constrained score by missense variants had much higher significance level than that by loss of function variants (*p*=1.46E-45 vs. 1.27E-06) although both predictors were also positively related to the counts of rare non-synonymous mutations. Interestingly, the interaction term between the coding region length and gene frequency score had negative effects on the mutation counts (−0.193±0.004) and had the most significance level according to their z-scores. The negative interaction suggested that frequency score has a smaller effect for large genes than for small genes. The genome may show local variation in mutation frequency. For a large gene, the local variation was averaged out so that the length mattered the most. For small genes the local information was important and this was captured by the frequency score. The varied tendency can be visualized among genes in three gene groups of coding region lengths (Figure 1c).

#### Alzheimer’s disease (AD)

Alzheimer’s disease is a progressive neurodegenerative disorder characterized by memory loss and cognitive impairments among elderly people. Multiple factors, including amyloid beta (Aβ) plaques, hyperphosphorylated tau tangles in the brain, synaptic dysfunction, neuronal loss, gut microbiota and neuroinflammatory, were recognized to be associated with the development and progression of AD ^39-41^. This dataset contained 246 ApoE ε4-negative AD patients and 172 healthy elderly controls sequenced at exomes. The detailed sample information can be seen in the original paper ^42^. RUNNER also showed approximately uniformly-distributed p-values at the most majority genes for the rare variant mutation burden test (Figure S2). However, the three alternative tests showed deflation in the small sample (Figure S2). The unweighted version of RUNNER detected a significant gene, CXCL16 (p=1.85E-6), which had 17 non-synonymous variants in the patients (See details in Table S3). It encodes a CX3C subgroup of chemokines and played a role in a wide range of diseases, including neuropathies and inflammatory diseases. Although no studies have established its genetic association with AD, there have been some indirect evidences. For instance, Rosito et al suggested that CXCL16 protected neurons from excitotoxic cell death in the CNS ^43^. Castro et al showed that CXCL16 modulated neurotransmitter release in hippocampal CA1 area^44^. RUNNER detected another gene as significant gene DPH1 (p=9.64E-7) (See details in Table S4). DPH1 encodes an enzyme involved in the biosynthesis of diphthamide. This gene had 19 non-synonymous or splicing variants in the patients. Most of the variants had large deleteriousness score which amplified the mutation counts in RUNNER. The estimated coefficients of six predictors of RUNNER in this dataset were similar to that in HSCR (Table S5). Except for the interaction predictor, all other predictors had positive coefficients. The interaction predictor also showed the highest statistical significance.

In contrast, the alternative methods detected no significant genes. The smallest p-value by CMC, KBAC and SKAT were 5.18E-4 (TULP3), 6.0E-4 (TBC1D1), and 1.16E-3 (ZNF296). None of these genes have been implicated in AD by previous studies. The CXCL16 was ranked by KBAC as the 2^nd^ gene as well. However, its association p-value (*p*= 7.0E-4) failed to survive the exome-wide threshold (2.5E-6).

## Discussion

In the present study, we proposed a novel approach, RUNNER, to evaluate baseline rare mutation burden across genes or regions in whole-genome or –exome sequencing studies. To our knowledge, this is the first statistical test for genetic mapping by baseline mutation burden across genes, which is carried out by a truncated negative binomial regression. We showed that RUNNER was much more powerful than three widely-used methods (CMC, SKAT and KBAC) for genetic association test at rare variants. The latter were built on mutation burden between cases and controls. We also demonstrated that RUNNER was much more robust to population structure than the widely-used methods. The enhanced power was reaffirmed by detecting more known or promising candidate susceptibility genes in real data analysis of Hirschsprung disease and Alzheimer’s disease. To facilitate application of this powerful approach, we have implemented RUNNER as one of analysis modules on our comprehensive software platform for high throughput sequencing data analysis, KGGSeq (http://grass.cgs.hku.hk/limx/kggseq/).

Why is RUNNER more powerful than a typical case-control association test? An intuitive explanation as follows: in a typical case-control test, the null hypothesis is that cases and controls have equal rare variant frequencies - the rare variant frequency in the entire population - estimated by the overall rare variant frequency in the pooled sample of cases and controls. However, in genomic regions that contain rare variants positively associated with disease, the use of a pooled estimate results in an inflation of the population rare variant frequency, and attenuates the apparent increase in rare variant frequencies in cases, leading to reduced statistical power. Instead of testing for deviations of case and control frequencies from a pooled frequency estimate, RUNNER tests for deviations from an estimate of population rare variant frequency obtained by statistical modelling of the rare variant frequency in the entire genome using available genomic and population data. For illustration, consider the hypothetical example of a disease with prevalence 0.01, and a genomic region where the frequency of rare variants is 0.001 per individual in controls, but 0.01 per individual in cases. The overall frequency of rare variants in the population is therefore 0.001*0.99+0.01*0.01= 0.00109. In a sample size 1000 cases and 1000 controls, the expected number of rare variants in the cases and controls would be 10 and 1, respectively. If these expected values were actually observed, then the pooled rare variant frequency would be 0.0055, while the Pearson chi-squared statistic would be approximately (10-5.5)^2^/5.5 + (1-5.5)^2^/5.5 = 7.37, producing a one-tailed p-value of 0.0033. However, if we were able to infer correctly, through statistical modelling, that the population frequency of rare variants in the genomic region is 0.00109, then the expected number of rare variants in 1000 cases would be 1.09, so that the probability of observing 10 or more variants assuming a Poisson distribution is 2.4E-7.

Besides the new strategy as illustrated in the results section, the enhanced power of RUNNER is also attributed to multiple methodological innovations. First, its iterative regression model based on the truncated negative binomial distribution exploits a baseline mutation burden comparison across genes or regions to detect disease risk genes. This is totally different from current mainstream analysis methods which are based on a case-control comparison. The comparison of baseline mutation burden across genes was seldom adopted for germline mutations analysis although it has been used for cancer somatic mutations analysis^20^. Probably, the major obstacle is the difficulty in modeling the distribution of mutation counts of rare variants. The proposed RUNNER provides a proper solution to the problem, which can be indirectly indicated by the approximated uniform distribution of the produced p-values among background genes. The improved power can further be explained by its higher sensitivity to increment of mutant alleles. RUNNER directly uses the increased allele numbers via the deviance residues to produce a test statistic instead of allele frequencies in conventional case-control association tests. In addition, the model also has an advantage of integrating prior weights at variants to further boost the power. As shown in Figure 3, the prior weights lead to 30% or more extra power while keeping the correct type 1 error.

The robustness of RUNNER to population stratification is also ascribed to the baseline mutation burden. The spurious association by conventional case-control studies is often caused a scenario that part or all cases are sampled from a population ancestrally different from that of controls. So the mutation burden between cases and controls only showed genetic difference between different populations. While the principle component analysis is often adopted to adjust for population stratification at common variants, it may be not effective for rare variant association analysis^14^. In contrast, the comparison resorts mutation burden across genes instead of that between cases and controls to detect susceptibility genes. It is much less insensitive to the population mixture. In our mimic population stratification where all controls were from an ancestrally different population (European vs. East Asian), RUNNER still showed no inflation of type 1 errors (Figure 3). This robustness relieves the difficulty in matching controls samples for rare variant-based association test. Theoretically, RUNNER also works for case-only samples as long as there are no private variants. Therefore, RUNNER had an advantage of tackling the long-lasting problem of population stratification issue in genetic association studies.

We selected six predictors under RUNNER model for the background mutation count of genes in the present study. Although the six predictors have led to accurate prediction and approximately uniform distribution of p-values, it does not exclude the usage of other predictors. For example, the sequencing coverage at a gene or region would be also a predictor. But it may be data-dependent. When sequencing depth is high enough, almost all genes have enough reads to call reliable genotypes. In this scenario, its impact on the prediction will be small and will have little effect on statistical evaluation (See an example in Figure S4). Moreover, it has been noted that the mutation rate tends to be lower in genes that are highly expressed ^45^, due to a process termed transcription-coupled repair ^46^. The mutation rate tends to be higher in regions with late DNA replication timing ^47^, probably due to the depletion of free nucleotides ^48^. Studies have suggested that mutation rates may be also associated with epi-genomic features ^49^. However, these features are cell-type and developmental stage dependent. In practice, it is often challenging to obtain a right cell-type at a right time for a disease. We tried to add the gene expression of multiple-cell types from GTEx ^50^ as predictors. It turned out there are generally insignificant (unpublished data). In fact, the frequency score predictor is generated at the same genome regions and may already have contained part of the impact of gene expression and replication timing.

The model of RUNNER is independent of mutation types. Due to availability of abundant data for validation, we focus on coding and splicing mutations in this paper. This does not preclude its potential extension to non-coding rare mutations. However, additional future work is needed for the extension to non-coding variants. As the evolution selection and genomic features of non-coding variants may be different from that of coding variants ^51^, different predictive factors may be needed for the mutation counts in non-coding regions. The biological interpretation of significant results is also more difficult for long non-coding genomic regions. A possible way is to analyze sub-non-coding regions separately, e.g., regulatory upstream regions and the first intronic regions. Besides, the functional prediction at non-coding variants is generally less advanced than that at coding variants. How the prior weights based on less accurate prediction at non-coding variants influence the power will also be an interesting future work. We performed a preliminary extension of the proposed framework for rare variants in upstream and downstream regions. Although only the gene-frequency score was used, the p-values had only slightly deviation from the uniform distribution (Figure S3). This result suggested the framework RUNNER is extendable for non-coding regions with more effective predictors.

Besides the systematic simulation study, the real data analysis further demonstrates effectiveness of RUNNER for genetic mapping of rare variants. In a proof of principle example, RUNNER successful “re-discovered” the known causal gene (p<2.5E-6), RET ^32; 33^, and prioritized another known gene EDNRB ^33; 34^ of Hirschsprung disease. These genes were ignored by the three widely-used alterative tests, CMC, SKAT and KBAC. In the Alzheimer’s disease dataset, RUNNER showed significant p-value at CXCL16. Although experiments are needed to validate the association, there have been multiple indirect evidences supporting this gene as a promising candidate gene of Alzheimer’s disease, including protection of neurons from excitotoxic cell death in the CNS ^43^ and regulation of neurotransmitter release in hippocampal CA1 area ^43^.

Two limitations should be noted in the present paper. First, the analysis framework does not directly consider covariates to adjust for confounding factors of diseases. For a disease with strong confounding factors, one should pre-adjust the case-control status or clinical phenotypes by a regression with the confounding factors and re-group the patients with the adjusted disease status or phenotypes. Second, RUNNER will not work in candidate gene analysis as it resorts to hundreds of genes to build a baseline model at a time. Therefore, the substantially enhanced power of RUNNER does not mean that it will be a replacement of the widely-used case-control tests for genetic mapping. However, it may work as an effective complementary analysis approach either when well-matched control samples are unavailable or when a genome-wide case-control tests are underpowered.

In summary, we proposed a novel and powerful statistical approach, RUNNER, to detect susceptibility genes with rare mutations. Its baseline mutation burden and advanced weighted truncated negative binomial regression may motivate follow-up methodological studies for genetic mapping from a new angle. As we demonstrated in the applications, the enhanced power of RUNNER may save multiple susceptibility genes missed by widely-used case-control association tests.

## Description of Supplemental Data

Figures S1–S4, Tables S1–S6, and Supplemental Methods

## Acknowledgements

This work was funded by National Natural Science Foundation of China (31771401 and 31970650), National Key R&D Program of China (2018YFC0910500 and 2016YFC0904300), Science and Technology Program of Guangzhou (201803010116), Theme-based Research scheme T12C-714/14-R.

## Web Resources

The website of KGGSeq: http://grass.cgs.hku.hk/limx/kggseq/

The website of gnomAD: https://gnomad.broadinstitute.org/

